# Powassan Virus NS5 Antagonizes TYK2-Mediated Immune Signaling Pathways

**DOI:** 10.64898/2026.09.07.749923

**Authors:** Mary O’Mara, Melissa Molho, Andrew P. Kurland, Zachary Walter, Ségolène Gracias, Vincent Caval, Nolwenn Jouvenet, Jeffrey R. Johnson, Holly Ramage

## Abstract

Powassan virus (POWV) is an emerging neurotropic tick-borne flavivirus, yet the mechanisms by which POWV evades host antiviral immunity remain poorly defined. Here, we identify multiple mechanisms of POWV innate immune antagonism with the viral polymerase NS5 protein as a central inhibitor of cytokine signaling. Both POWV lineages potently inhibited type I interferon (IFN) signaling, and NS5 expression suppressed signaling and downstream interferon-stimulated gene expression. Affinity purification-mass spectrometry identified the host kinase TYK2 as a conserved NS5 interactor. POWV NS5 binds the TYK2 kinase domain through a discrete interface within the RNA-dependent RNA polymerase (RdRp) region between catalytic motifs B and C and inhibits TYK2 phosphorylation. Disruption of this interface abrogated TYK2 binding and reduced NS5-mediated IFN antagonism, while revealing additional TYK2-independent mechanisms of immune suppression. POWV NS5 also inhibited TYK2-dependent IFN-λ and IL-12 signaling, demonstrating that its immune antagonism extends beyond Type I IFN. Together, these findings identify TYK2 as a central target of POWV immune evasion and implicate the variable RdRp B-C region as an interface for flavivirus-host interactions. More broadly, our results reveal how POWV can coordinately suppress multiple antiviral cytokine pathways and provide insight into mechanisms that may shape tick-borne flavivirus host adaptation and pathogenesis.

## INTRODUCTION

Powassan virus (POWV) is an emerging tick-borne neurotropic flavivirus endemic to North America and parts of Asia^1^. Like other members of the *Orthoflavivirus* genus, POWV possesses a ∼11 kb positive-sense RNA genome that encodes a single polyprotein which is cleaved by host and viral proteases into three structural proteins (C, prM, and E) which comprise the viral particle, and seven nonstructural proteins (NS1, NS2a, NS2b, NS3, NS4a, NS4b, and NS5) which are involved in several steps of the virus life cycle and host immune modulation^2,3^.

There are two genetic lineages of POWV in circulation in North America, which share 93% amino acid identity and are serologically indistinguishable: Lineage I, and Lineage II, also called deer tick virus (DTV)^4–9^. In nature, POWV is maintained in a sylvatic cycle between small mammal reservoirs and tick vectors, with transmission to humans occurring incidentally through the bite of an infected tick^10^. Most POWV infections in humans are asymptomatic or self-limiting, characterized by mild symptoms including fever or headache; however, spread of POWV into the central nervous system can cause encephalitis, seizures, and death^11^. POWV encephalitis has a 10% case fatality rate, with half of survivors experiencing long-term neurological sequelae such as weakness, tremors, and paralysis^12^. While POWV infection in humans is relatively rare, there has been a steady increase in reported cases within the United States and Canada over the last several years^13^. Further, surveillance studies suggest that up to 5% of ticks in some regions of the United States test positive for POWV^14^ and the increasing prevalence of POWV infection is likely to be exacerbated by the expanding host range of the tick vectors^15^. Despite this growing public health concern, no approved vaccines or antiviral therapies are currently available, underscoring the need for a deeper understanding of POWV pathogenesis and host-virus interactions.

Host defense against viral infection relies on the rapid detection of pathogen-associated molecular patterns by innate immune sensors, including the cytosolic RNA sensor RIG-I, which recognizes viral RNA and replication intermediates^16,17^. Activation of these pathways triggers production of type I interferons (IFNs), key antiviral cytokines that signal through the type I interferon receptor (IFNAR) to activate the JAK-STAT pathway and induce expression of hundreds of interferon-stimulated genes (ISGs)^18^. These ISGs establish an antiviral state that restricts viral replication and spread. Consequently, successful viral infection depends on the ability to evade or suppress Type I IFN responses^16^.

While the general strategy to antagonize the production and downstream signaling of Type I IFNs is conserved, the specific mechanisms employed by related flaviviruses are quite diverse^19^. These strategies have been described for a subset of flaviviruses but have not been determined across the *Orthoflavivirus* genus. For tick-borne flaviviruses, viral NS5 protein binding to TYK2 has been proposed as a major mechanism for Type I IFN antagonism for tick-borne encephalitis virus (TBEV) and Louping Ill virus (LIV), but additional tick-borne flaviviruses such as POWV have not yet been characterized^20^. To address this gap, we sought to determine whether POWV can antagonize Type I IFN signaling and to define the mechanisms mediating innate immune evasion.

Here, we show that POWV potently antagonizes Type I IFN signaling through the viral polymerase NS5, which suppresses STAT1/2 phosphorylation and downstream ISG expression. Mechanistically, POWV NS5 interacts with the tyrosine kinase domain of the host kinase TYK2, impairing its phosphorylation and thereby reducing IFNAR signaling strength. We further map this interaction to a discrete region within the NS5 RNA-dependent RNA polymerase domain and demonstrate that disruption of NS5-TYK2 binding attenuates IFN antagonism. Beyond Type I IFN signaling, NS5-mediated targeting of TYK2 broadly impairs additional TYK2-dependent cytokine pathways, including IFN-λ and IL-12 signaling. Together, these findings identify TYK2 as a central target of POWV immune evasion and reveal a mechanism by which tick-borne flaviviruses suppress multiple arms of the host antiviral response.

## RESULTS

### POWV NS5 antagonizes the host Type I IFN response

To determine whether POWV can antagonize Type I IFN signaling, we infected HMC3 microglial cells with Lineage II POWV Spooner (POWV SP) virus, stimulated cells with IFN-β, and measured the total and phosphorylated forms of the Type I IFN signaling protein STAT2, by western blotting. While total STAT2 protein levels were unchanged, POWV SP-infected cells had significantly reduced levels of phosphorylated STAT2 (**Fig. 1A**). Phosphorylated STAT1 and STAT2 interact with IRF9 to form the transcription factor ISGF3, which translocates to the nucleus and binds to the IFN-stimulated response element (ISRE), driving the expression of interferon-stimulated genes (ISGs)^21,22^. Next, we measured ISG expression in mock-infected or POWV SP-infected cells with and without IFN-β stimulation. We found that POWV-infected cells had significantly lower transcription of ISGs (*Ifit1*, *Mx1*, *Socs1*, *Rig-I)* than mock-infected cells upon Type I IFN stimulation (**Fig. 1B, Fig. S1A**). These data demonstrate that POWV antagonizes Type I IFN signaling and downstream ISG expression in HMC3 cells.

**Figure 1.**
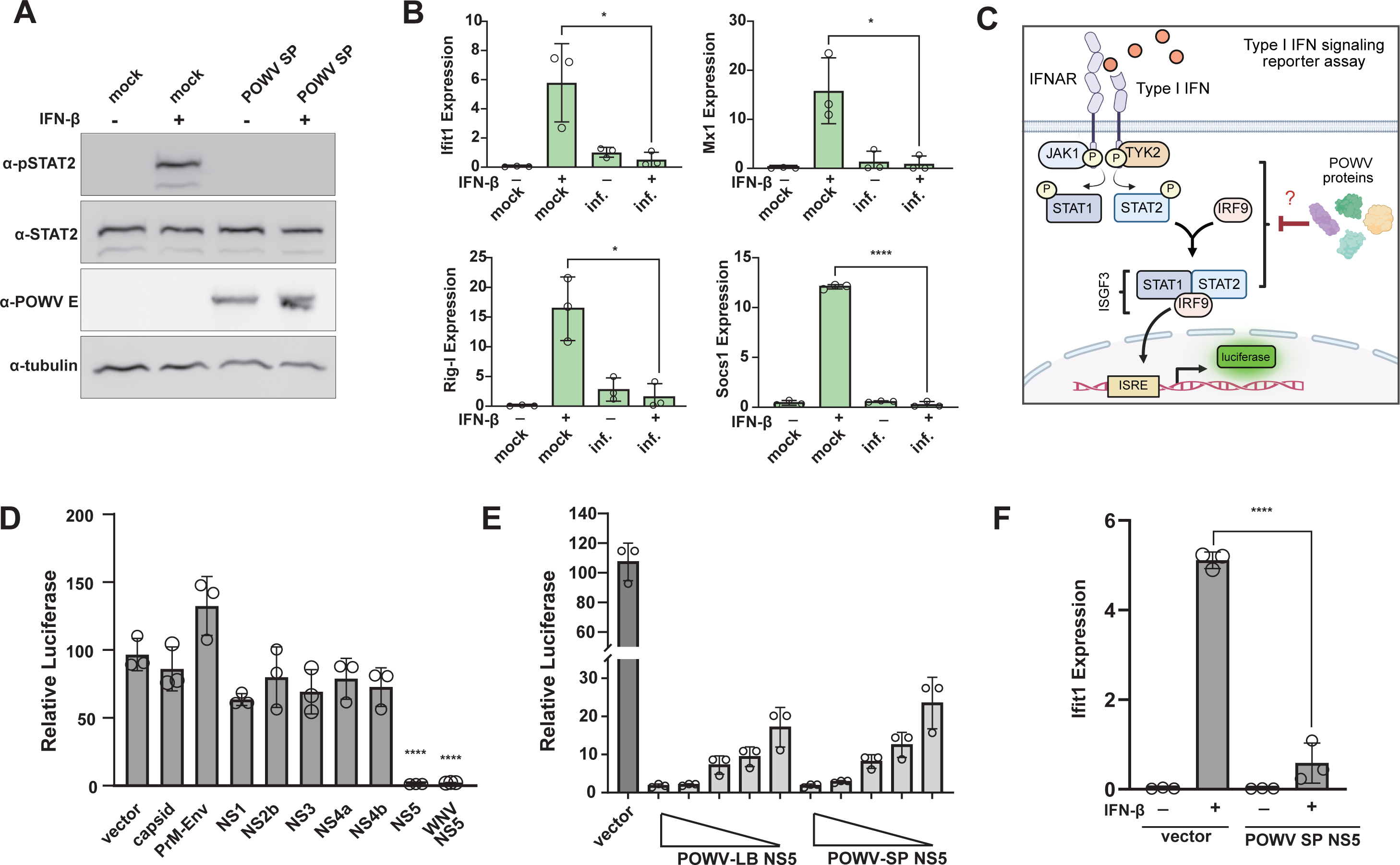
POWV NS5 antagonizes the host Type I IFN response. (**A**) HMC3 microglial cells were infected with POWV SP (MOI 0.1 for 48hr), stimulated with IFN-β (20ng/mL) for 20min, then subjected to western blotting with the indicated antibodies (*n*=3 independent experiments; representative images shown). (**B**) HMC3 cells were infected with POWV SP (MOI 0.1 for 48hr), stimulated with IFN-β (20ng/mL) for 3hr, then subjected to RNA extraction and qRT-PCR to measure the indicated mRNAs, normalized to 18S RNA (*n*=3 independent experiments; Student’s unpaired *t*-test, two-tailed). (**C**) Schematic depicting luciferase reporter assay for ISRE-driven gene expression. (**D**, **E**) HEK293T cells were co-transfected with luciferase reporter plasmids and individual POWV LB proteins, stimulated with IFN-β (5ng/mL) for 18hr, then lysates were assayed for luciferase activity (*n*=3 independent experiments; ordinary one-way ANOVA with Šidák’s multiple comparisons test). (**F**) HMC3 stably expressing POWV NS5 or empty vector control were stimulated with IFN-β (20ng/mL) for 3hr, then subjected to RNA extraction and qRT-PCR for *Ifit1* mRNA normalized to 18S RNA (*n*=3 independent experiments; Student’s unpaired *t*-test, two-tailed). For all data, the mean ± SD is shown. Significance is indicated by ^∗^*p* < 0.05, or ^∗∗∗∗^*p* < 0.0001.

Related flaviviruses have been shown to antagonize Type I IFN signaling through multiple distinct mechanisms, including via interactions between viral and host proteins^16,20,23–25^. To identify the mechanism by which POWV antagonizes this pathway, we first investigated whether expression of any individual POWV protein was sufficient to antagonize Type I IFN signaling. We cloned each of the ten viral proteins (Lineage I, LB) into a Strep-tagged vector and confirmed protein expression of each construct (N-terminal tag for capsid, C-terminal for all other proteins, **Fig. S1B**). We co-transfected each of these vectors with a reporter construct encoding luciferase under the control of the interferon-sensitive response element (ISRE) promoter (**Fig. 1C**). We observed that POWV NS5 expression strongly inhibited luciferase activity downstream of Type I IFN stimulation (**Fig. 1D**). This inhibition was similar in scale to that of WNV NS5, which has been previously demonstrated to antagonize Type I IFN signaling^26^. Furthermore, NS5 proteins from both the Lineage I (LB) and Lineage II (SP) POWV strains antagonized ISRE-driven luciferase production in a dose-dependent manner (**Fig. 1E, Fig. S1C**). We also observed that expression of POWV NS5 was sufficient to antagonize transcription of an ISG (*Ifit1)* in response to Type I IFN stimulation (**Fig. 1F**). To determine if individual POWV proteins can also antagonize production of Type I IFN, we co-transfected each of the POWV LB protein constructs with a construct expressing GST-RIG-I(2CARD)^27^ and a reporter construct encoding luciferase under the control of the IFN-β promoter. We observed inhibition of IFN-β production upon expression of POWV LB NS1, similar to the inhibition observed upon expression of DENV-2 NS2a, which has been shown to antagonize the production of IFN-β^28^ (**Fig. S1D**). Together, these results demonstrate that POWV antagonizes Type I IFN through multiple mechanisms and that POWV NS5 is the primary viral protein inhibiting signaling downstream of IFNAR.

### POWV NS5 interacts with host antiviral factors

We reasoned that POWV antagonism of Type I IFN signaling may be mediated by interactions between NS5 with one or more host proteins. To identify NS5-interacting host factors, we expressed POWV LB NS5 and POWV SP NS5 with a C-terminal Strep tag in both HEK293T and HMC3 microglia cells and performed affinity purification. Expression of NS5 was confirmed by both silver stain and western blotting (**Fig. S2A, B**). Next, we identified NS5-interacting host proteins by a data-independent acquisition mass spectrometry (DIA-MS) approach, yielding quantification of 2,935 protein groups (**Fig. 2A, S2C, Table S1**). Principal component analysis of DIA-MS data showed strong separation by cell type in PC1, while PC2 separated cells transfected with NS5 constructs compared to vector control (**Fig. S2D)**. To compare proteins enriched in NS5 purifications compared to negative control, data were scored by the MSstats and SAINTq algorithms. Protein groups were filtered for log2fold-change (NS5/Control) > 1, adjusted p-value < 0.05, SAINTq BFDR < 0.01, and the number of peptides > 1. We identified a total of 118 protein groups passing these criteria (**Table S1, S2E**). Functional term enrichment analysis of the proteins passing these filtering criteria revealed a strong enrichment for splicing factors (U2 snRNP; U2-type pre-spliceosome assembly) in HEK293T cells, as with TBEV NS5^29^, and an enrichment for the phosphorylase b kinase complex in HMC3 cells (**Fig. 2B**). Next, we generated a network view of NS5-interacting proteins by integrating the BioGRID multivalidated protein-protein interaction data set (**Fig. 2C**)^30^. We identified protein interaction subnetworks that passed our interaction scoring threshold for both POWV strains and in both cell lines: a PML-ATRX-SMN-GEMI4 complex comprised of proteins localized to defined subnuclear locations, and a SETD2-IWS1 complex involved in coupling transcription elongation and chromatin modification^31^. Finally, we highlighted POWV NS5-interacting proteins that were also found to interact with tick-borne encephalitis virus (TBEV) and Louping ill virus (LIV) NS5 proteins^20^. Notably, we identified an interaction with the non-receptor tyrosine kinase protein TYK2 with both POWV strains, in both cell lines, that has also been described to interact with TBEV and LIV NS5, both in cells expressing NS5 and in cells infected with TBEV^20,29^.

**Figure 2.**
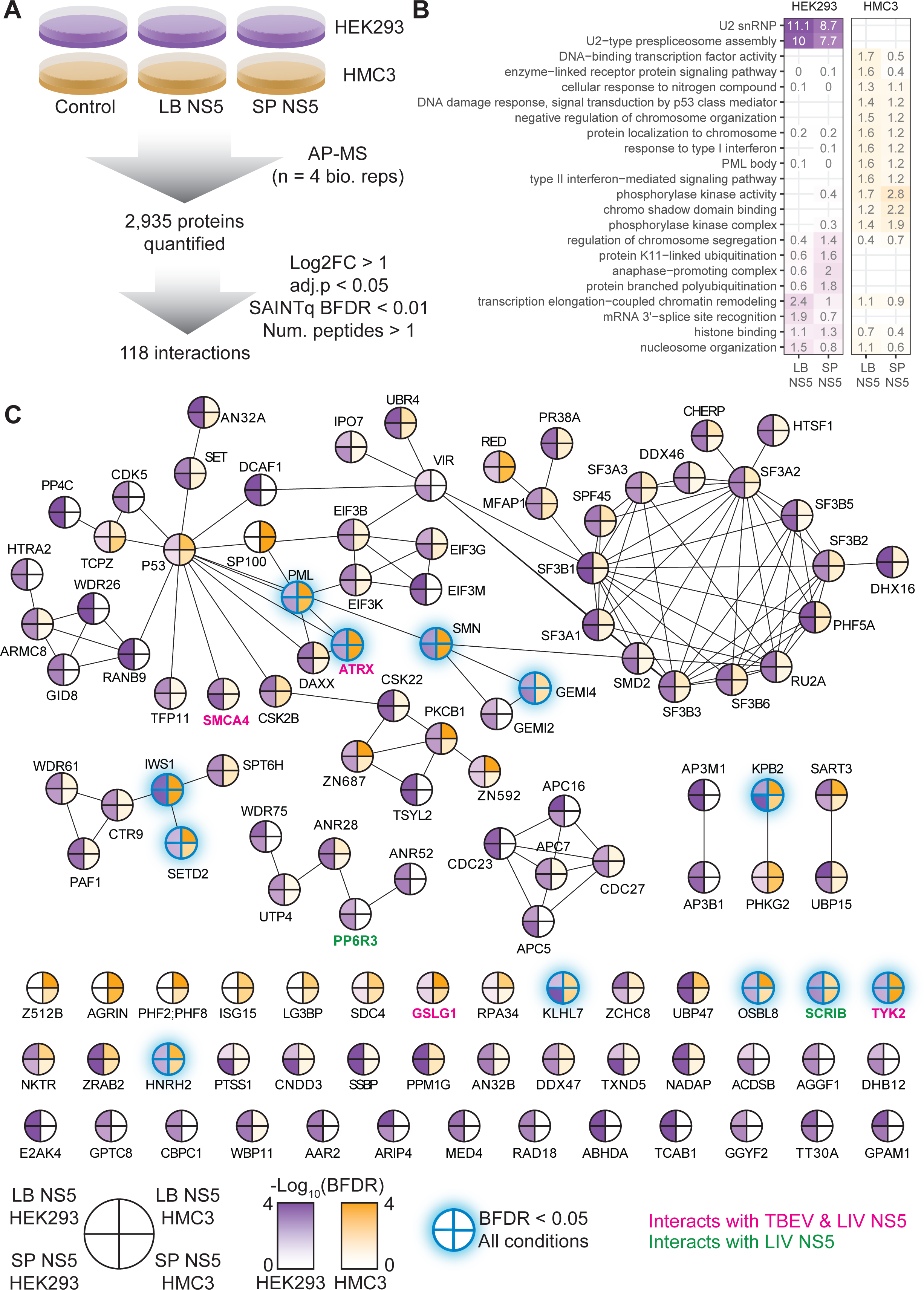
**POWV NS5 interacts with multiple host proteins**. (**A**) Strep-tagged POWV NS5 (from LB or SP) was expressed and affinity-purified from HEK293T and HMC3 cells in biological quadruplicate. Material purified from cells expressing NS5 and untransfected cells was analyzed by DIA-MS and interactions scored using the MSstats and SAINTq algorithms. (**B**) Heatmap of -log_10_(adjusted *p*-value) of enrichment of gene ontology terms using the gProfiler2 package in R^82^. (**C**) Interaction network for proteins defined to interact with POWV NS5 based on criteria in (A). Node fill indicates the -log_10_(SAINTq BFDR) for each POWV strain purified from each cell line; nodes highlighted in blue with a blue shadow passed interaction criteria for both POWV strains purified from both cell types; edges indicate protein interactions in the BioGRID multivalidated data set; labels in green and pink indicate interactions overlapping with TBEV and LIV NS5 proteins.

### POWV NS5 interacts with the enzymatic domain of TYK2 and antagonizes its phosphorylation

To validate the POWV NS5-TYK2 interaction, we performed co-immunoprecipitation experiments with tagged POWV NS5 constructs. We observed that POWV LB and SP NS5 co-purified with both overexpressed V5-tagged TYK2 as well as with endogenous TYK2, confirming this interaction, while WNV NS5 did not interact with TYK2 (**Fig. 3A-B**). We next sought to identify the regions of TYK2 and POWV NS5 needed for their interaction. Similar to other Janus kinases (JAKs), TYK2 consists of four major domains. The N-terminus of TYK2 contains the 4.1, Exrin, Radixin, Moesin (FERM) and Src homology 2 (SH2) domains, which mediate an interaction with the cytoplasmic portions of the Type I interferon (IFNAR) receptor. The C-terminus contains the enzymatically active tyrosine kinase (TK) domain capable of phosphorylation of other Janus kinases, such as JAK1, as well as the homologous but enzymatically inactive kinase-like domain (KL)^32,33^. We generated V5-tagged domain truncation mutants of TYK2, then co-transfected these constructs along with Strep-tagged full-length POWV NS5 in HEK293T cells (**Fig 3C**). Only constructs expressing the tyrosine kinase (TK) domain co-purified with POWV NS5, suggesting that the interaction of TYK2 with POWV NS5 is mediated by the TYK2 TK domain (**Fig. 3D**).

**Figure 3.**
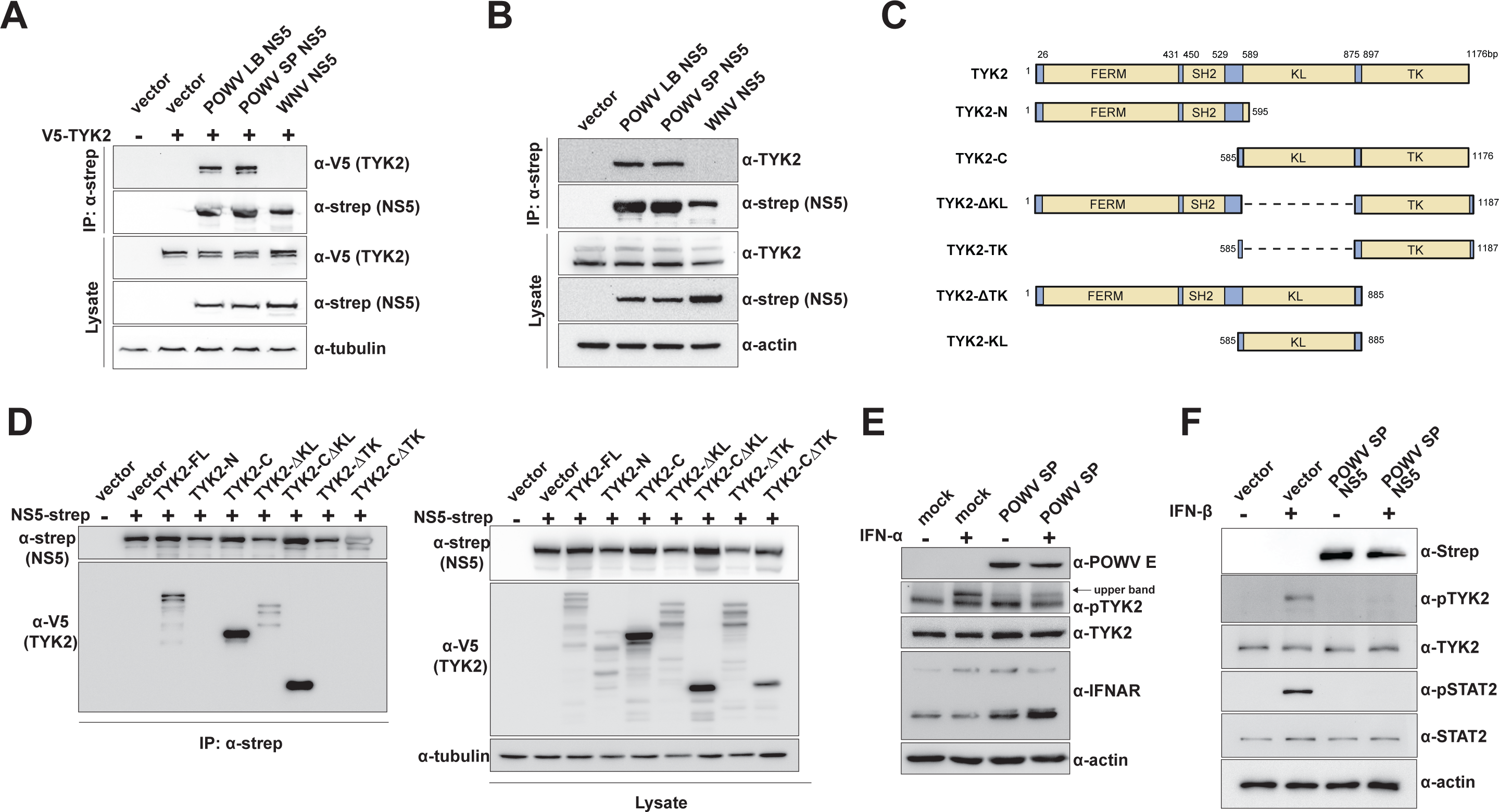
POWV NS5 interacts with the enzymatic domain of TYK2 and antagonizes its phosphorylation. (**A**) HEK293T cells were transfected with V5-tagged TYK2, Strep-tagged POWV LB NS5 or POWV SP NS5, Strep-tagged WNV NS5, or an empty vector control. Lysates were subjected to affinity purification followed by western blotting with the indicated antibodies (*n*=3 independent experiments; representative images shown). (**B**) HEK293T cells were transfected with Strep-tagged POWV NS5 or controls. Lysates were subjected to affinity purification followed by western blotting with the indicated antibodies (*n*=3 independent experiments; representative images shown). (**C**) Schematic depicting TYK2-V5 truncation mutants. (**D**) HEK293T cells were transfected with V5-TYK2 truncation constructs and Strep-tagged POWV-NS5. Lysates were subjected to affinity purification followed by western blotting with the indicated antibodies (*n*=3 independent experiments; representative images shown). (**E**) Vero E6 cells were infected with POWV SP (MOI 0.1 for 48hr), stimulated with IFN-α (3000 IU/mL) for 15min, then lysed and probed with the indicated antibodies (*n*=3 independent experiments; representative images shown). (**F**) HMC3 cells stably expressing POWV NS5 or controls were stimulated with IFN-β (20 ng/mL) for 20min, then lysed and probed with the indicated antibodies (*n*=3 independent experiments; representative images shown).

Since phosphorylation of TYK2 allows for signal transduction through IFNAR, we hypothesized that the interaction between POWV NS5 with the enzymatic domain of TYK2 might antagonize Type I IFN-mediated TYK2 phosphorylation. To test this, we mock-treated or POWV SP-infected Vero E6 cells, stimulated with IFN-α, and performed western blotting on the cell lysates. While total IFNAR and TYK2 protein levels were similar between mock and infected cells, POWV-infected cells had reduced Type I IFN-induced TYK2 phosphorylation (**Fig. 3E**). Furthermore, HMC3 microglia cells stably expressing POWV LB NS5 or POWV SP NS5 also had impaired TYK2 phosphorylation in response to Type I IFN as measured by western blot (**Fig. 3F**). Together, these data suggest that POWV NS5 antagonizes phosphorylation of TYK2.

### POWV NS5 interaction with TYK2 reduces Type I IFN signaling

In order to test the requirement for the TYK2 interaction on NS5 antagonism of Type I IFN, we first determined the region of POWV NS5 responsible for interacting with TYK2. We co-transfected HEK293T cells with expression constructs encoding TYK2-V5 and either of the two enzymatic domains of NS5: the N-terminal methyltransferase (MTase) domain or the C-terminal RNA-dependent RNA polymerase domain (RdRp) (**Fig. 4A**). Co-immunoprecipitation and western blotting revealed that the RdRp-only construct, but not the MTase-only construct, co-purified with TYK2 (**Fig. 4B**). Furthermore, the POWV NS5 RdRp domain, but not the MTase domain, was sufficient to impair ISRE-driven reporter gene expression in response to Type I IFN stimulation (**Fig. 4C**).

**Figure 4.**
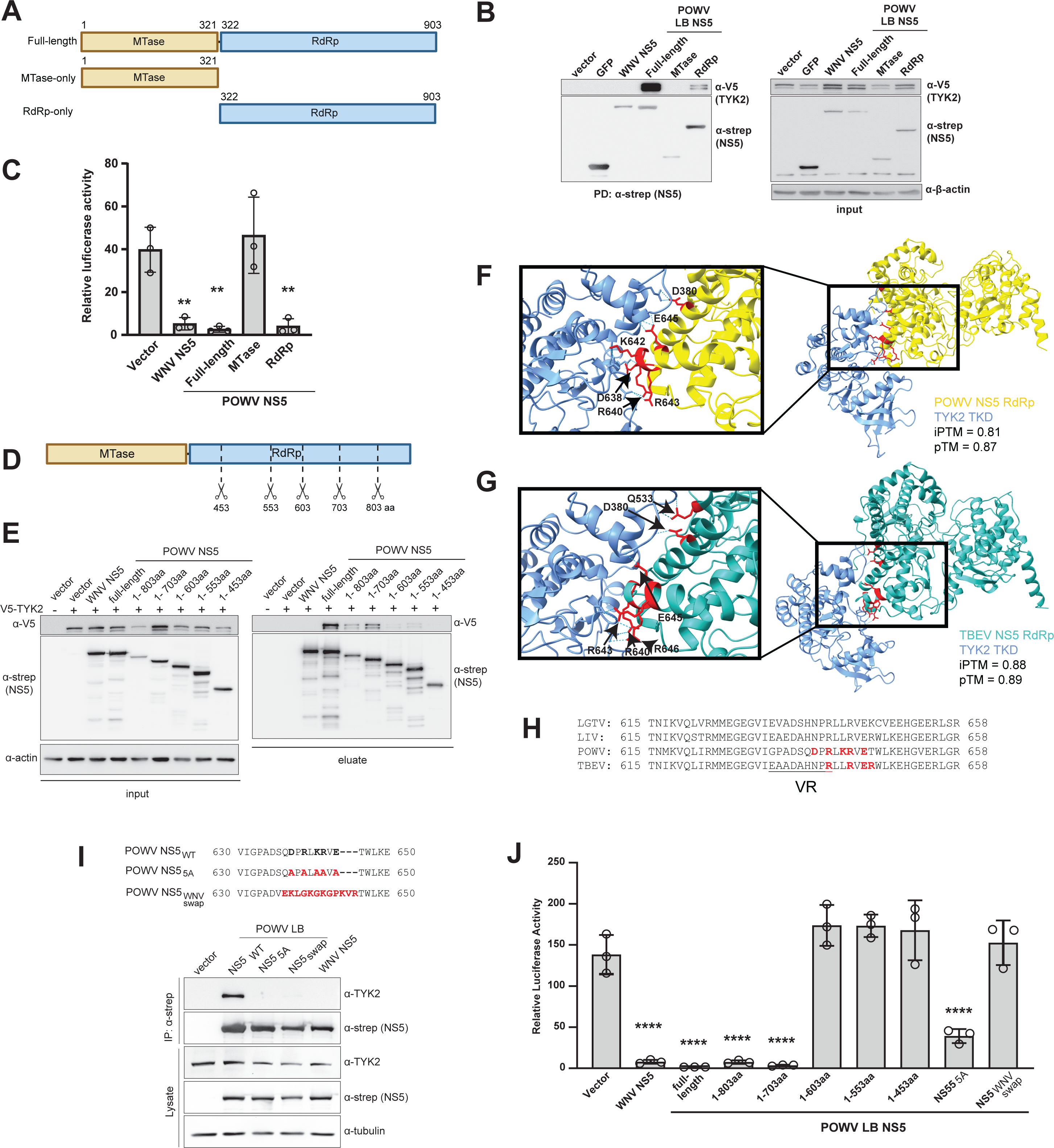
POWV NS5 interaction with TYK2 contributes to Type I IFN antagonism. (**A**) Schematic depicting Strep-POWV LB NS5 domain truncation mutants. (**B**) HEK293T cells were co-transfected with TYK2-V5 and Strep-POWV LB NS5 domain truncation mutants or controls. Lysates were subjected to affinity purification followed by western blotting with the indicated antibodies (*n*=3 independent experiments; representative images shown). (**C**) HEK293T cells were co-transfected with luciferase reporter plasmids and Strep-POWV LB NS5 domain truncation mutants, stimulated with IFN-β (5ng/mL) for 18hr, then lysates were assayed for luciferase activity (*n*=3 independent experiments; ordinary one-way ANOVA with Šidák’s multiple comparisons test). (**D**) Schematic depicting Strep-POWV LB NS5 RdRp C-terminal truncations. (**E**) HEK293T cells were co-transfected with TYK2-V5 and Strep-POWV LB NS5 full-length, C-terminal truncation mutants, or WNV NS5 and empty vector controls. Lysates were subjected to affinity purification followed by western blotting with the indicated antibodies (*n*=3 independent experiments; representative images shown). (**F, G**) AlphaFold3 prediction of interaction between POWV NS5 RdRp (**F**) or TBEV NS5 RdRp (**G**) and the TYK2 TK domain with interacting residues indicated. Accuracy prediction scores are shown out of a 0 (low) to 1 (high) scale; iPTM, interface predicted template modeling; pTM, predicted template modeling. (**H**) Alignment of residues for LGTV, LIV, POWV, and TBEV in region predicted to interact with TYK2. Interacting residues predicted by AlphaFold in red. (**I**) HEK293T cells were transfected with Strep-POWV LB NS5 wild-type and mutant constructs (mutations indicated in red), WNV NS5, or empty vector controls and subjected to affinity purification and western blotting with the indicated antibodies (*n*=3 independent experiments; representative images shown). (**J**) HEK293T cells were transfected with Strep-POWV LB NS5 full-length, or the indicated mutant constructs, WNV NS5 or an empty vector control and stimulated with IFN-β (5ng/mL) for 18hr. Lysates were assayed for luciferase activity (*n*=3 independent experiments; ordinary one-way ANOVA with Šidák’s multiple comparisons test). For all data, the mean ± SD is shown. Significance is indicated by ^∗∗^*p* < 0.005, or ^∗∗∗∗^*p* < 0.0001.

To more finely map the region of interaction, we generated constructs encoding the NS5 MTase domain with successive C-terminal 100aa truncations of the RdRp domain (**Fig. 4D**). Co-IP and ISRE-luciferase assays with these constructs revealed that NS5 residues 603-703 within the RdRp domain are necessary for the interaction of POWV NS5 with TYK2 (**Fig. 4E**), as well as for the antagonism of ISRE-driven gene expression (**Fig. 4J**). We next performed *in silico* analysis using AlphaFold3 to predict interacting residues between the TK domain of TYK2 and the interaction region in the POWV NS5 (RdRp)^34^. This yielded high-confidence interaction scores involving a total of six charged residues in the POWV NS5 RdRp domain (**Fig. 4F**). Five of those six predicted interacting residues fell within the region of the NS5 RdRp (603-703aa) that we previously determined was required for the interaction with TYK2 (**Fig. 4E**). POWV is closely related to tick-borne encephalitis virus (TBEV). Since it has recently been observed that the TBEV NS5 RdRp domain also interacts with TYK2^20^, we also used AlphaFold3 to predict the TYK2-interacting residues of TBEV NS5 (**Fig. 4G**). The predicted TYK2-interacting residues for both POWV NS5 and TBEV NS5 are near the variable region of the NS5 B-C loop. Furthermore, three of the five POWV residues predicted to mediate the TYK2 interaction are conserved with TBEV and also predicted to mediate the TBEV NS5 interaction with TYK2 (**Fig. 4H**).

Next, we generated two POWV NS5 constructs bearing mutations at these predicted TYK2-binding residues. We mutated each of the five predicted interacting residues in this region to alanine (POWV LB NS5_5A_), or swapped a 12 amino acid region encompassing these five residues with that of WNV NS5, which does not interact with TYK2 (POWV LB NS5_WNV_) (**Fig. 4I**). We co-transfected these constructs into HEK293T cells with TYK2-V5 and performed affinity purification and western blotting. We observed that both the POWV LB NS5_5A_ and POWV LB NS5_WNV_ constructs failed to co-immunoprecipitate with TYK2, suggesting that these five residues in the POWV NS5 RdRp are critical for the interaction with TYK2 (**Fig. 4I**). Finally, we incorporated these mutants into our ISRE-luciferase reporter assay and found that both constructs had impaired antagonism of ISRE-driven gene expression, as compared to wild-type POWV NS5 (**Fig. 4J**). Interestingly however, the POWV LB NS5_5A_ construct was still able to significantly inhibit Type I IFN signaling, suggesting that additional interactions between POWV NS5 and other host factors may mediate additional mechanisms of antagonism of Type I IFN signaling, independent of the TYK2 interaction (**Fig. 4J**).

### POWV NS5 antagonizes additional TYK2-dependent pathways

In addition to its role in Type I IFN signaling, TYK2 is also required for other immune signaling pathways, including Type III IFN (IFN-λ), IL-12, and IL-23 signaling **(Fig. 5A**)^35^. Given that POWV NS5 antagonizes TYK2 to impair Type I IFN signaling, we wanted to determine if NS5 can also antagonize these additional pathways. Production and sensing of IFN-λ is a key host antiviral defense mechanism in barrier tissues including the skin and gut which, along with Type I IFN signaling, results in the production of ISGs. HepG2 hepatocytes have previously been shown to be responsive to IFN-λ, and we observed that pre-treatment of HepG2 cells with IFN-λ exerted a strong protective effect from infection with POWV SP (**Fig. 5B**)^36^. To test whether POWV NS5 antagonizes IFN-λ signaling, we generated HepG2 hepatocyte and HaCaT keratinocyte cell lines stably expressing POWV LB or SP NS5. We observed that in both cell lines, expression of POWV NS5 impaired transcription of the ISG IFIT1 in response to treatment with IFN-λ. (**Fig. 5C-D, Fig. S3A-B**). As there may be crosstalk between the Type I and Type III signaling pathways, we directly tested whether signaling downstream of IFN-λ is inhibited in POWV-infected cells. We infected HepG2 cells with POWV SP for 48 hours and treated uninfected or infected cells with IFN-β or IFN-λ for 20 minutes. We observed a more robust response to IFN-β, as compared to IFN-λ, consistent with what has been previously reported^37,38^ (**Fig. 5E**). As before (Fig. 1A), we observed that POWV-infected HepG2 cells inhibited STAT2 phosphorylation in response to IFN-β stimulation (**Fig. 5E**). In addition, we also found that POWV infection antagonizes STAT2 phosphorylation downstream of IFN-λ. Together, these results show that POWV NS5 inhibits Type III IFN signaling.

**Figure 5.**
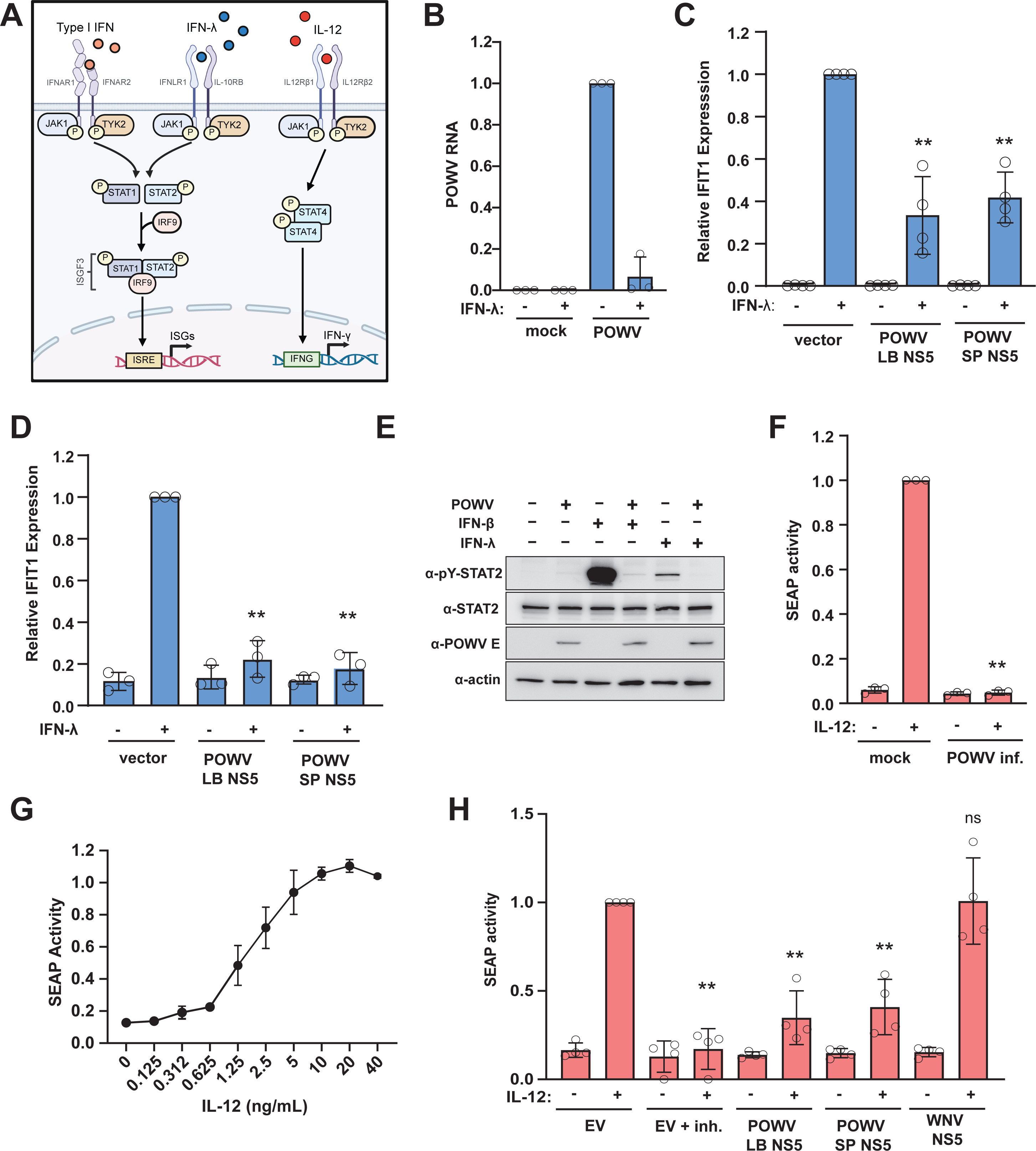
POWV NS5 antagonizes IFN-λ and IL-12 signaling. (**A**) Schematic depicting TYK2 involvement in IFN-λ and IL-12 signaling. (**B**) HepG2 hepatocyte cells were stimulated with IFN-λ_1_ (50ng/mL) for 24hr, infected with POWV SP (MOI 1 for 48hr), then subjected to RNA extraction and qRT-PCR for POWV SP NS5 RNA normalized to 18S RNA (*n*=3 independent experiments). (**C, D**) HepG2 hepatoma cells (**C**) or HaCaT keratinocyte cells (**D**) stably expressing POWV NS5 or empty vector controls left unstimulated or stimulated with IFN-λ_1_ (50ng/mL) for 24hr, then subjected to RNA extraction and qRT-PCR. *Ifit1* expression was normalized to 18S RNA (*n*=3 independent experiments; ordinary one-way ANOVA with Šidák’s multiple comparisons test). (**E**) HMC3 cells were infected with POWV SP (MOI 0.1 for 48h), left unstimulated or stimulated with 50ng/mL IFN-λ_1_ or 5ng/ml IFN-β for 20min, then lysates were subjected to western blotting. (**F**) HEK-Blue IL-12 cells were infected with POWV SP (MOI 1 for 48h), left unstimulated or stimulated with IL-12 (100ng/mL) for 24hr, and SEAP reporter enzyme activity measured in the supernatant (n=3 independent experiments; Mann-Whitney test). (**G**) HEK-Blue IL-12 cells stably expressing an empty vector control were stimulated with increasing concentrations of IL-12 for 24h and the reporter enzyme activity was measured in the supernatant (n=3 independent experiments). (**H**) HEK-Blue IL-12 cells stably expressing WNV NS5, POWV LB NS5, POWV SP NS5 or an empty vector control were left unstimulated or stimulated with IL-12 (100ng/mL) for 24hr, and the reporter enzyme activity was measured in the supernatant (n=3 independent experiments; Brown-Forsythe ANOVA with Dunnett’s T3 multiple comparisons test). For all data, the mean ± SD is shown. Significance is indicated by ^∗∗^*p* < 0.005.

TYK2 kinase activity is also implicated in IL-12 signaling, which is a critical cytokine in the development of antiviral Th1 CD4+ T cells (**Fig. 5A**). To test whether POWV can antagonize the effects of IL-12 stimulation, we used a reporter cell line (HEK-Blue IL-12 reporter cells), which stably express the IL-12 receptor as well as the scaffolding molecule STAT4. Upon IL-12 treatment, an alkaline phosphatase reporter enzyme (SEAP) is secreted into the cell culture supernatant. We mock-infected or POWV-infected reporter cells, stimulated the cells with IL-12 or left them unstimulated, and measured activity of secreted reporter SEAP. We found that POWV infection robustly limited the response of the reporter cells to IL-12 (**Fig. 5F, S3C**). Next, we tested whether the NS5 protein of POWV mediates antagonism of IL-12 signaling. We used lentiviruses to generate HEK-Blue IL-12 cells stably expressing POWV LB, POWV SP, or WNV NS5 and confirmed that the modified reporter cell line was still responsive to IL-12 stimulation (**Fig. 5G**). Next, we stimulated the NS5-expressing reporter cells with IL-12 and measured SEAP activity in the supernatant. We observed that the cells expressing POWV LB or SP NS5, but not WNV NS5, had impaired IL-12 signaling (**Fig. 5H, S3D**). As a positive control, we included the FDA-approved TYK2 inhibitor deucravacitinib and found a similar level of inhibition^39^ (**Fig. 5H**). These data show that POWV NS5 can antagonize IL-12 signaling. Together, our results show that the POWV NS5 inhibits signaling downstream of multiple TYK2-dependent immune pathways.

## DISCUSSION

Innate immune signaling, including Type I IFN, is critical in controlling viral infection; therefore, viruses must circumvent this restriction to establish a productive infection in the host. Viral antagonism of host antiviral responses can occur at multiple points along innate immune activation pathways, including the sensing of viral PAMPs, production of immunostimulatory molecules, and propagation of signal transduction cascades resulting in the production of antiviral effectors^16,40^. Individual flaviviruses have been observed to target a wide array of different host factors, and many target multiple factors in the same pathway, such as both the production of Type I IFN and signaling downstream of IFNAR^41^. In particular, signaling at and downstream of IFNAR is often targeted by the flaviviral NS5 protein^20,23,24,42–48^. However, the strategies used by POWV to subvert innate immune signaling have not been defined. To address this gap, we used orthogonal cell-based and biochemical approaches to identify POWV-host interactions that mediate immune evasion.

First, we demonstrated that both lineages of POWV antagonize the Type I IFN response (Fig 1A, B). Using an ISRE-luciferase reporter assay, we show that this antagonism is mediated by the POWV NS5 protein, consistent with the mechanisms identified in other tick- and mosquito-borne flaviviruses (Fig. 1D-F). As flaviviruses often act to inhibit the production of Type I IFN, we also tested the effect of expression of each POWV protein on an IFN-β driven luciferase assay and found that the POWV NS1 protein inhibits the production of IFN-β after stimulation with a RIG-I (2CARD) construct^27,49^ (Fig. S1D). It has been previously shown the WNV NS1 protein inhibits activation of the IFN-β promoter by promoting RIG-I degradation^50^. Further experimentation is needed to determine whether POWV NS1 acts in a similar manner or instead inhibits a distinct step downstream of pattern recognition receptor (PRR) sensing.

To determine how POWV NS5 may antagonize Type I IFN signaling, we performed affinity purification of POWV NS5 followed by mass spectrometry to identify interacting host proteins. To identify a set of high-confidence interactors, we performed these studies in biological quadruplicate using NS5 strains representative of both POWV lineages (POWV LB and POWV SP) and in two different cell lines (HEK293T and HMC3 microglia) (Fig. 2). These data revealed 12 host proteins that were identified in all data sets. Further, we identified six host proteins that were also found to interact with TBEV and/or LIV NS5^20^. Among these interactors was TYK2, a Janus-activated kinase which undergoes activating phosphorylation upon stimulation of IFNAR. This phosphorylation leads to further signal propagation, ultimately resulting in the production of antiviral interferon stimulated genes (ISGs). Importantly, TYK2 was found in all conditions in our dataset and has been shown to interact with both TBEV and LIV NS5, in both NS5-expressing cells and in TBEV-infected cells^20,29^ (Fig. 2).

We confirmed the POWV-TYK2 interaction by affinity purification and western blotting and demonstrated that the interaction interface maps to the RdRp domain of POWV NS5 and the TK domain of TYK2 (Fig 2A-D, 4A-B). We used *in silico* predictions and found five specific residues within the POWV NS5 RdRp that are predicted to interact with the TYK2 TK domain. These residues are within or directly adjacent to the variable region between catalytic motifs B and C (region B-C) in the RdRp^51^ (Fig. 4F, H). Our modeling as well as a previous report show that the residues mediating the interaction between the TBEV NS5 RdRp and TYK2 are also located in this region, with three predicted TYK2-interacting residues shared between POWV and TBEV NS5 (R640, R643, E645) (Fig. 4H)^20,51^. Significant sequence and structure variations have been observed in the RdRp region B-C in flaviviruses, and this region has been shown to mediate innate immune evasion in WNV and LGTV^26,52^. The sequence plasticity of the RdRp B-C region, together with its role in innate immune antagonism, suggests that this region may function as an evolutionary interface for virus-host interactions, enabling flaviviruses to adapt to species-specific antiviral pressures.

Importantly, we validated the importance of the residues within this region for binding TYK2 by generating two POWV NS5 mutants: a construct in which all five predicted interacting residues were mutated to alanine (POWV-N55_5A_), and one in which the 12aa region encompassing these five residues was swapped with the homologous region of WNV NS5 (POWV-NS5_WNV_). Our *in vitro* binding experiments show that both of these mutants abrogate interaction with TYK2 (Fig. 4I). Additional studies will establish the contribution of individual amino acids in NS5 for maintaining the interaction with TYK2.

Crucially, we demonstrated that POWV NS5 mutants which fail to interact with TYK2 exhibit reduced antagonism of innate immune signaling pathways (Fig. 4J). However, while both constructs failed to co-immunoprecipitate with TYK2, only the POWV-NS5_WNV_ construct fully lost the ability to antagonize Type I IFN signaling, while the POWV-N55_5A_ construct still retained robust inhibitory activity (Fig. 4J). These data suggest that there may be additional mechanisms by which POWV NS5 can antagonize Type I IFN signaling, independent of the interaction with TYK2. Other flavivirus NS5 proteins can inhibit signaling through a reduction in cell surface expression of IFNAR, or through the targeted degradation of downstream signaling proteins (e.g. STAT1, STAT2)^23,24,45–48^. However, we did not observe reduced expression of total STAT2 or IFNAR in POWV-infected cells, suggesting a distinct mechanism by which POWV antagonizes Type I IFN signaling (Fig. 1A, 3E). Further studies to determine the additional components or signaling steps that are impacted by expression of the POWV-N55_5A_ mutant, and a comparative study to identify host interaction partners of the POWV-N55_5A_ as compared to WT NS5 may shed light on these other mechanisms of innate immune antagonism.

While the impact of Type I IFN signaling antagonism in the context of flavivirus infection is well-described, studies on the antagonism of other cytokine signaling pathways by flaviviruses are more limited. The interaction and inhibition of TYK2 by tick-borne flavivirus NS5 led us to examine antagonism of other TYK2-dependent antiviral pathways likely to be important for controlling POWV infection. TYK2 is a key component in IFN-λ signaling, which also mediates ISG expression^37,53^. The kinetics of the antiviral response to IFN-λ is generally slower than the response to Type I IFN^37^, and the production and recognition of Type III IFN is largely limited to subtypes of immune cells as well as epithelial cells in barrier surfaces as the gut, skin, placenta, and blood-brain barrier (BBB)^54,55^. However, studies have demonstrated an important role for IFN-λ in maintaining the integrity of the blood-brain barrier to prevent viral neuroinvasion in mosquito-borne flavivirus infection^56,57^.

Upregulation of Type III IFN is observed in cells infected with POWV^58^. More recently, a protective role of IFN-λ *in vitro* was shown upon pre-treatment for TBEV and POWV infection, and our data are consistent with these findings (Fig. 5B)^59^. We show that both POWV LB and SP NS5 can inhibit downstream signaling and ISG expression after stimulation with IFN-λ (Fig. 5C, D). This suggests that the interaction between POWV NS5 and TYK2 may be important for antagonizing innate immune responses at barrier surfaces. IFN-λ has been shown to play a critical role in protection at the maternal-fetal interface, restricting vertical transmission of ZIKV^60,61^. Interestingly, it was shown that POWV can also infect the fetus of pregnant mice and cause fetal demise, and that POWV can infect human fetal tissue^62^. This suggests the possibility of vertical transmission of POWV and a potential protective role of IFN-λ in this context.

TYK2 mediates signaling via other proinflammatory cytokines, including IL-12 and IL-23. IL-12 is synthesized early in viral infection during dendritic cell activation and maturation, bridging the innate and adaptive immune response. IL-12 activates macrophages and cytotoxic T cells, while IL-23 promotes viral clearance by a Th17 response that recruits and activates neutrophils to the site of infection^63–65^. Macrophages and dendritic cells are early targets of POWV infection^66^. It is well-established that these cells produce IL-12 and IL-23 to drive the immune response to viral infection, and macrophages and dendritic cells can also respond to these cytokines in an autocrine manner^67–71^.

Several studies have shown that flavivirus infection can impair the maturation of dendritic cells, inhibiting their downstream immune functions and promoting viral infection^72–76^. Moreover, antagonism of IL-12 production has been observed during infection of dendritic cells with the related tick-borne flavivirus Langat virus (LGTV)^77^. Using IL-12 reporter cells, we show that both POWV infection and expression of the POWV NS5 protein can antagonize IL-12 signaling (Fig. 5F-H). Together, these data suggest that tick-borne flaviviruses may inhibit both the production and the downstream signaling of IL-12, pointing to a critical role for this cytokine in restricting infection. Interestingly, our data show that expression of WNV NS5 does not inhibit IL-12 signaling, which may indicate a tick-borne flavivirus-specific mechanism of immune evasion (Fig. 5H). However, it is possible that WNV and other flaviviruses antagonize IL-12 signaling via a mechanism that does not require the viral NS5 protein. Further studies are required to determine whether the antagonism of IL-12 signaling is mediated by the POWV NS5-TYK2 interaction and whether this interaction can also antagonize IL-23 signaling during infection.

Together, our findings reveal that POWV employs multiple strategies to counter host antiviral immunity and that the NS5 protein is an antagonist of multiple antiviral cytokine signaling pathways. By targeting TYK2, a shared signaling component downstream of multiple cytokines, POWV may disrupt distinct arms of the antiviral response across different cell types, tissues, and stages of infection. These findings broaden the significance of flavivirus NS5-mediated immune evasion beyond the well-characterized antagonism of Type I IFN. Furthermore, the identification of the variable RdRp B-C region as a host factor binding interface suggests that this region may be an evolutionary hotspot through which flaviviruses acquire host- and virus-specific immune evasion strategies. Future studies defining the relative contributions of Type I IFN, IFN-λ, IL-12, and potentially IL-23 antagonism during infection, and determining whether these mechanisms are conserved across tick-borne flaviviruses, will provide important insight into how immune evasion shapes flavivirus host adaptation and pathogenesis.

## MATERIALS AND METHODS

### Cells and viruses

HEK293T (ATCC CRL-1573), VeroE6 (ATCC CRL-1586) and HaCaT (Fisher Scientific NC0309203) cells were grown under standard conditions in Dulbecco’s Modified Eagle Medium (Corning 10-017-CM) supplemented with 10% fetal bovine serum (R & D Systems, Cat S12450) and 2mM L-alanyl-L-glutamine dipeptide (GlutaMax, Gibco). HMC3 (ATCC CRL-3304), HepG2 (ATCC HB-8065), and BHK-21 (ATCC CCL-10) cells were grown under standard conditions in Eagle’s Minimum Essential Medium (Corning 10-009-CV) supplemented with 10% fetal bovine serum. HEK-Blue IL-12 cells (Invivogen hkb-il12) were cultured according to manufacturer’s instructions. Cell cultures were routinely tested for mycoplasma contamination via PCR (MycoStrip Mycoplasma Detection Kit, Invivogen rep-mys-100) and maintained for less than 30 passages. Catalog numbers for each cell line are listed in the supplementary table below. Powassan Virus LB (M794) was obtained from the UTMB Arbovirus Reference Collection, virus Pool Number TVP23083. The POWV SP isolate was obtained from UTMB Arbovirus Reference Collection, virus pool number TVP 20467. Viral titers were determined in BHK-21 cells by TCID_50_ assay.

### Plasmids and generation of mutants

To clone POWV genes, viral RNA was isolated from infected Vero E6 cells and complementary cDNA was amplified using Transcriptor First-Strand cDNA Synthesis Kit (Roche) and a 3’ reverse POWV LB oligo, following the manufacturer’s instructions. POWV genes were cloned into pCDNA4_TO with a C-terminal 2X Strep II affinity tag for all proteins except capsid which was cloned into pCDNA4_TO with a N-terminal 2X Strep II affinity tag using CloneAmp HiFi PCR premix and In-Fusion Cloning (Takara Bio). A list of primers used for cloning, mutagenesis, and sequencing is provided in **Table S2**. All inserts were verified using Sanger sequencing followed by full-length Nanopore sequencing (Plasmidsaurus). The V5-tagged TYK2 constructs used in this paper are described in^20^. The lentiviral packaging plasmids pMD2.5 (Addgene 12259) and psPAX2 (Addgene 12260) and lentiviral expression plasmid pCDH-Str-Ii (Addgene 65313) were gifted from Andrew South. The pCDH-Str-Ii plasmid was mutagenized to remove Ii and add C-terminally strep-tagged POWV LB NS5, POWV SP NS5, and WNV NS5 using CloneAmp HiFi PCR premix and In-Fusion Cloning (Takara Bio). Plasmids used in this study are listed in **Table S3**.

### Transfections

1.5e5 HMC3 cells were seeded in 6-well plates, then transfected the next day with 1-2ug DNA depending on experiment at a volumetric ratio of 1:3 with X-tremeGENE™ 9 DNA Transfection Reagent (Roche 06365787001). HEK293T cells were seeded at 5e5 per well in 6-well plates and transfected with 1-2ug DNA depending on experiment at a volumetric ratio of 1:3 with Polyjet In Vitro DNA Transfection Reagent (SignaGen SL100688) according to manufacturer’s instructions.

### Luciferase assays

1.25e5 HEK293T cells were seeded in 24-well plates and transfected the next day with 10-200 ng of POWV DNA, 200ng pGL4.45 ISRE-firefly luciferase, and 10ng pNL1.1 NanoLuc Renilla luciferase plasmids (Promega). After 24-48hrs, cells were stimulated with 5ng/mL IFN-β for 18hrs. For the IFN-β luciferase assay, cells were seeded as above and transfected with 10-200 ng of POWV DNA, 200ng IFN-beta_pGL3 (Addgene 102597), 10ng pNL1.1 NanoLuc Renilla luciferase and 50 ng of RIG-I 2CARD plasmid and analyzed after 48hr. Cells were lysed in passive lysis buffer and freeze-thawed, then lysates were assayed using the Nano-Glo dual-luciferase reporter assay (Promega N1610) following the manufacturer’s instructions to record activity in lysates of both firefly luciferase (ISRE-driven) and Renilla luciferase (CMV-driven, to control for any changes in global transcription). Absorbance was measured at all wavelengths (firefly luciferase) and at 460nm (Renilla luciferase) on a Molecular Devices SpectraMax iD3 plate reader. Measurements were subtracted from the signal of the untransfected control cells to account for background luminescence. Values are reported as relative luciferase activity are the ratios of firefly luciferase over Renilla luciferase activity.

### Lentiviral transduction

HEK293T cells seeded in 6-well plates were transfected with XtremeGENE9 following manufacturer’s instructions with 0.7ug pMD2.G (Addgene 12259), 1ug psPAX2 (Addgene 12260), and 1ug of pCDH_Strep_POWV LB NS5, pCDH_Strep_POWV SP NS5, or pCDH_Strep_WNV NS5. After 48 hours, lentiviral supernatant was harvested, passed through a sterile 0.45um syringe filter, aliquoted, and frozen at -80C. HMC3, HepG2, and HaCaT cells were seeded in 6-well plates and transduced with 50ul of lentiviral supernatant. After 48 hours, cells were split and underwent selection with puromycin (Invivogen ant-pr-1) at 1ug/mL (HMC3) or 10ug/mL (HepG2 and HaCaT). After one week of selection, stocks of the surviving cells were aliquoted and frozen. Cells were maintained in puromycin following the second passage after recovery from thawing. Puromycin was omitted from media when seeding cells for functional assays.

### Immunoprecipitation

Cells were rinsed with 1x PBS, lysed in ice-cold IP buffer (150mM NaCl, 50mM Tris pH 7.4, 1mM EDTA) with 0.5% (v/v) Nonident NP-40 substitute (Sigma 74385), freeze-thawed, and homogenized by passing multiple times through a 23G needle and syringe. Lysates were centrifuged at 20min/4C/20,000xg and whole cell lysates set aside. The remaining cleared sample was added to 40ul of Strep-Tactin® Sepharose® 50% suspension (IBA 2-1201-002) and incubated rotating end-over-end at 4C overnight. The resin was washed four times with 500ul of IP buffer with 0.5% (v/v) Nonident NP-40 substitute or 1x RIPA buffer (50mM Tris-HCl pH 8.0, 150mM NaCl, 1% (v/v) Nonidet NP-40 substitute, 0.5% (w/v) sodium deoxycholate, 0.1% (w/v) SDS), then incubated with 50ul of 5mM biotin (IBA 6-6325-001) in IP buffer without NP-40 substitute rotating end- over-end at 4C for 1hr, centrifuged, and beads collected with mass spectrometry analysis and supernatant collected for western blotting.

### Sample preparation for mass spectrometry

Beads were washed once and resuspended with 40µl of 2M urea, 50 mM Tris pH 8.0, and 1mM DTT, and incubated for 30min at 37°C with shaking at 700 rpm. Iodoacetamide (Sigma-Aldrich) was added to 3mM final concentration and incubated at room temperature for 45min in the dark with shaking. Digestion was performed by adding 3mM DTT and 750ng of sequencing-grade modified trypsin (Promega) and incubating at 37°C with shaking overnight. Finally, supernatants were collected to fresh tubes and an additional 500ng of sequencing-grade modified trypsin was added and incubated for 2hr at 37°C with shaking. Digested samples were desalted using C18 solid phase extraction columns (C18 BioPurSPN, Nest Group) according to the manufacturer’s instructions. Desalted samples were vacuum centrifuged to dryness and resuspended in 0.1% formic acid for mass spectrometry analysis.

### Data acquisition for mass spectrometry

All samples were analyzed on an Orbitrap Eclipse mass spectrometry system equipped with an Easy nLC 1200 ultra-high pressure liquid chromatography system interfaced via a Nanospray Flex nanoelectrospray source (Thermo Fisher Scientific). Samples were injected onto a fritted fused silica capillary (30cm × 75μm inner diameter with a 15μm tip, CoAnn Technologies) packed with ReprosilPur C18-AQ 1.9 μm particles (Dr. Maisch GmbH). Buffer A consisted of 0.1% formic acid in water, and buffer B consisted of 0.1% formic acid in 80% acetonitrile. Peptides were separated by an organic gradient from 5% to 35% mobile buffer B over 60min, followed by an increase to 100% B over 10min at a flow rate of 300nL/min. Analytical columns were equilibrated with 3μL of buffer A. To build a spectral library, samples from each set of biological replicates were pooled and acquired in a data-dependent manner. Data-dependent acquisition (DDA) was performed by acquiring a full scan over a m/z range of 375-1025 in the Orbitrap at 120,000 resolving power (@200m/z) with a normalized AGC target of 100%, an RF lens setting of 30%, and an instrument-controlled ion injection time. Dynamic exclusion was set to 30 seconds, with a 10 p.p.m. exclusion width setting. Peptides with charge states 2-6 were selected for MS/MS interrogation using higher energy collisional dissociation (HCD) with a normalized HCD collision energy of 28%, with 3 seconds of MS/MS scans per cycle. Data-independent analysis (DIA) was performed on all individual samples. A full scan was collected at 60,000 resolving power over a scan range of 390-1010m/z, an instrument controlled AGC target, an RF lens setting of 30%, and an instrument controlled maximum injection time, followed by DIA scans using 8m/z isolation windows over 400-1000m/z at a normalized HCD collision energy of 28%.

The mass spectrometry proteomics data have been deposited to the ProteomeXchange Consortium via the PRIDE partner repository with the dataset identifier PXD082916^78^.

### Data processing for mass spectrometry

The Spectronaut algorithm was used to build spectral libraries from DDA data, identify peptides/proteins, and extract intensity information from DIA data^79^. DDA data were searched against the Homo sapiens UniProt reference proteome (downloaded on August 23 2023) and POWV sequences. False discovery rates were estimated using a decoy database strategy. All data were filtered to achieve a false discovery rate of 0.01 for peptide-spectrum matches, peptide identifications, and protein identifications. Search parameters included a fixed modification for carbamidomethyl cysteine and variable modifications for N-terminal protein acetylation and methionine oxidation. All other search parameters were Biognosys factory defaults. Statistical analysis of proteomics data was conducted utilizing the MSstats package in R^80^. All data were normalized by equalizing median intensities, the summary method was Tukey’s median polish, and the maximum quantile for deciding censored missing values was 0.999. Proteomics data were subsequently analyzed with the SAINTq algorithm^81^. For SAINTq analysis, analysis was performed at the fragment level using the normalized peak area reported by Spectronaut.

### Structural modeling

Sequences of the POWV NS5 RdRp domain and TYK2 tyrosine kinase domains were used as inputs for AlphaFold3 modeling. The top-scoring model was imported into ChimeraX (version 1.12) for H-bind and salt-bridge prediction using default parameters (radius of 0.075 Å, distance tolerance of 0.4 Å, and angle tolerance of 20°).

### Western blotting

Cells were washed with PBS and lysed in RIPA buffer supplemented with protease inhibitor cocktail (Thermo Scientific 78438) as well as phosphatase inhibitor cocktail when probing of phosphorylated proteins (Thermo Scientific 78442). Lysates were freeze-thawed and clarified by centrifugation at 20,000xg for 15min at 4C. Clarified lysates were transferred to new tubes, diluted with 6x Laemmli reducing sample buffer (BioWorld 10570021), and boiled for 5min. Protein samples were separated on 6%, 8%, or 10% SDS-PAGE gels and transferred to 0.45um nitrocellulose membrane (BioRad 1620115) using 20% methanol/0.05% SDS transfer buffer on ice with constant voltage at 100V/90min in a Mini-Protean chamber (BioRad). After blocking with 5% (w/v) non-fat dry milk in TBS-T for 1hr at room temperature, membranes were incubated with the indicated primary antibodies overnight at 4C, washed three times with TBS-T, and incubated with HRP-conjugated secondary antibodies for one hour at room temperature. For detection of pY-TYK2, the blocking, primary, and secondary antibody incubation steps were all performed overnight at 4C; the addition of HALT 1x protease/phosphatase inhibitor (ThermoScientific 78442) added to the blocking buffer and antibody dilutions during these incubations greatly enhanced the signal:noise ratio. Blots were washed three times with TBS-T, then incubated with ECL (Cytiva RPN3244) or SuperSignal West Femto Maximum Sensitivity Substrate (Thermo Scientific 34096) and imaged on an Amersham Imager 680 or iBright FL1500 (Invitrogen). Raw image files were converted from .fcz to .tif format using AlphaView (v3.5.0.927, Bio-Techne) and contrast, brightness, and alignment were adjusted in Photoshop (v. 24.7.5, Adobe). A list of primary and secondary antibodies is provided in **Table S4**.

### RNA isolation and quantitative RT-PCR

Total cellular RNA was isolated using Trizol (ThermoFisher Scientific) and purified and DNase-treated using Direct-zol RNA Purification Kit (Zymo Research R2051) per manufacturer’s instructions. Complementary DNA (cDNA) was synthesized using the Transcriptor High Fidelity cDNA synthesis kit (Roche 5081955001) using 1μg of input RNA with using random hexamer primers (ThermoFisher Scientific 48190011) with M-MLV reverse transcriptase (ThermoFisher Scientific 28025021) in a total volume of 20μl. The cDNA reactions were diluted 1:5 and 20μl of each diluted sample was used to make a pooled reference. The pooled reference was used for subsequent 10-fold dilutions to generate a standard curve for all targets being measured. cDNA reactions were further diluted 1:5 (1:25 total dilution) and SYBR green reactions contained 5μl of 2x PowerUp SYBR green (ThermoFisher Scientific A25743), 5μl of diluted cDNA, 5pmol of both forward and reverse primers, analyzed by qPCR and the relative abundance of each target was calculated using the standard curve. The relative values for each transcript were normalized to a control RNA (18S rRNA) and compared between experimental conditions. Primer sequences for qRT-PCR are listed in **Table S5**.

### SEAP assays

Supernatant from HEK-Blue IL-12 cells was pipetted from plates and centrifuged at 300xg for 5min to remove any residual cells. Supernatants were moved to new tubes and assayed following the manufacturer’s instructions (Invivogen). A 96-well plate with 180ul of QuantiBlue Solution and 20ul of sample per well was incubated for 1hr at 37C/5% CO2, then absorbance at 650nm was recorded using a SpectraMax iD3 plate reader (Molecular Devices).

### Quantification and statistical analysis

Statistical analyses were performed using R (proteomics) or Graphpad Prism. Details for specific experiments can be found in figure legends. Statistical tests include Student’s *t*-test (unpaired) or one-way ANOVA with Bonferroni correction for multiple comparisons. For each experiment, *n* is indicated in the figure legend and is defined as one biological replicate. Data are presented as mean ± SD or SE as indicated in figure legends. Significance was defined as an adjusted *p*-value (corrected for multiple comparisons if necessary) of below 0.05.

## Acknowledgements

We thank Dr. Michaela Gack for reagents used in this study. This work was supported by NIH/NIAID grant R01AI143850 to HR, Pennsylvania Department of Health Research Formula Research Grant SAP #4100095617 to HR, NIH/NIAID grants R21AI187731 and R01AI124690 to JRJ, and the NIH T32GM144302 grant to ZW and MO. Schematics were generated using BioRender.

## Supplemental Figure Titles and Legends

**Supplemental Figure 1 (Related to Figure 1).**
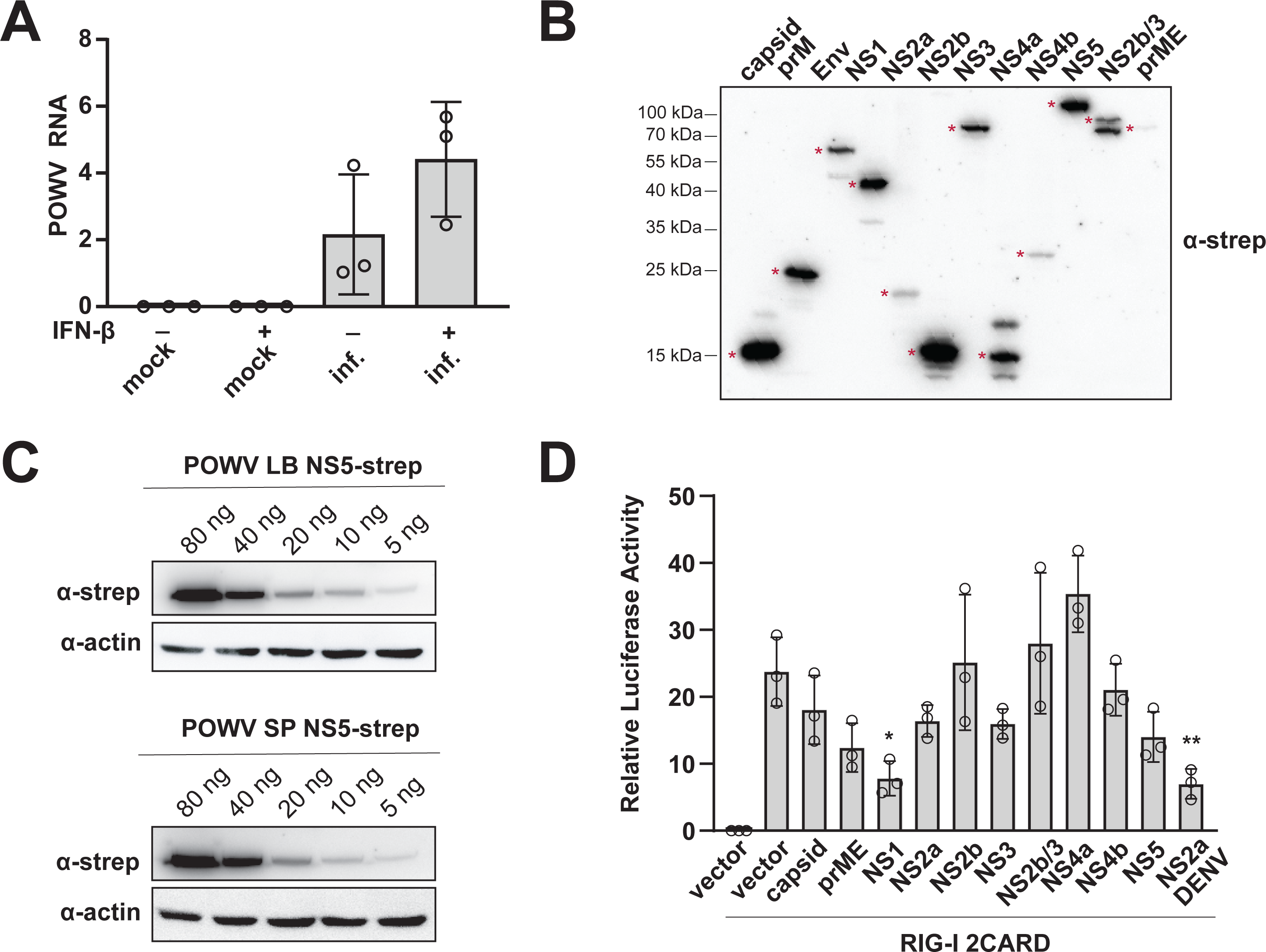
(**A**) qRT-PCR for POWV SP RNA of infected cells in Fig. 1B. (**B**) HEK293T cells were transfected with Strep-tagged POWV LB proteins and lysates were probed with anti-Strep antibody to validate expression. Bands corresponding to correct size are indicated with a red asterisk (*). (**C**) HEK293T cells were transfected with Strep-tagged POWV LB or SP NS5 at decreasing concentrations for ISRE luciferase assays (Fig. 1E) and lysates were probed with anti-Strep antibody to validate expression. (**D**) HEK293T cells were transfected with Strep-tagged POWV proteins and lysates analyzed using a IFN-β luciferase reporter assay (*n*=3 independent experiments; ordinary one-way ANOVA with Šidák’s multiple comparisons test). The mean ± SD is shown. Significance is indicated by ^∗∗^*p* < 0.005.

**Supplemental Figure 2 (Related to Figure 2).**
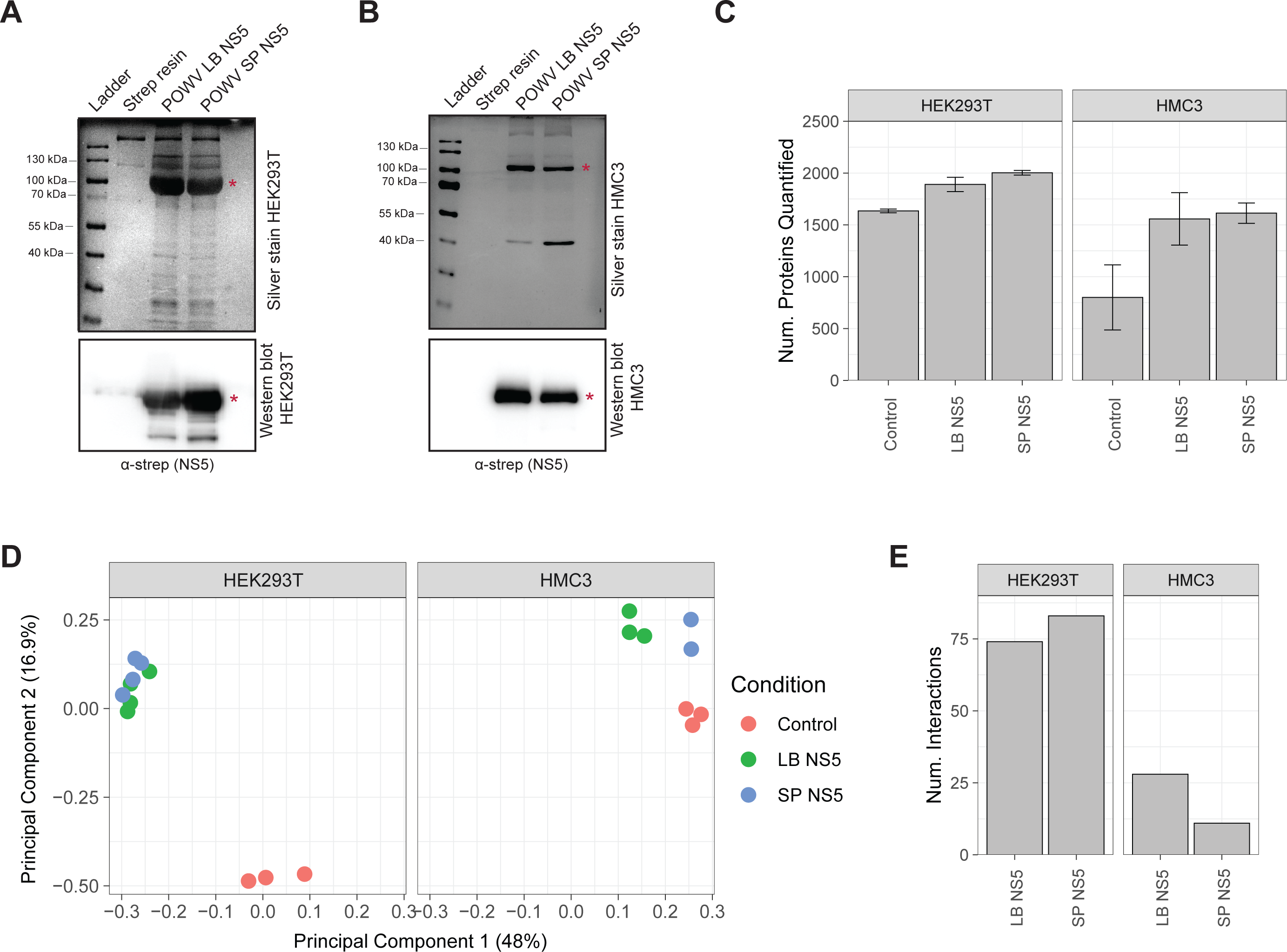
Lysates from HEK293T cells (**A**) and HMC3 cells (**B**) were transfected with Strep-tagged POWV LB NS5 and POWV SP NS5, separated by SDS-PAGE and visualized with silver staining (upper panel) and analyzed via western blot with an anti-Strep antibody (lower panel). Proteins of correct size are indicated with a red asterisk (*). (**C**) Number of proteins quantified in each sample condition. (**D**) Principal component analysis of all sample conditions. (**E**) Number of protein interactors identified in each sample condition.

**Supplemental Figure 3 (Related to Figure 5).**
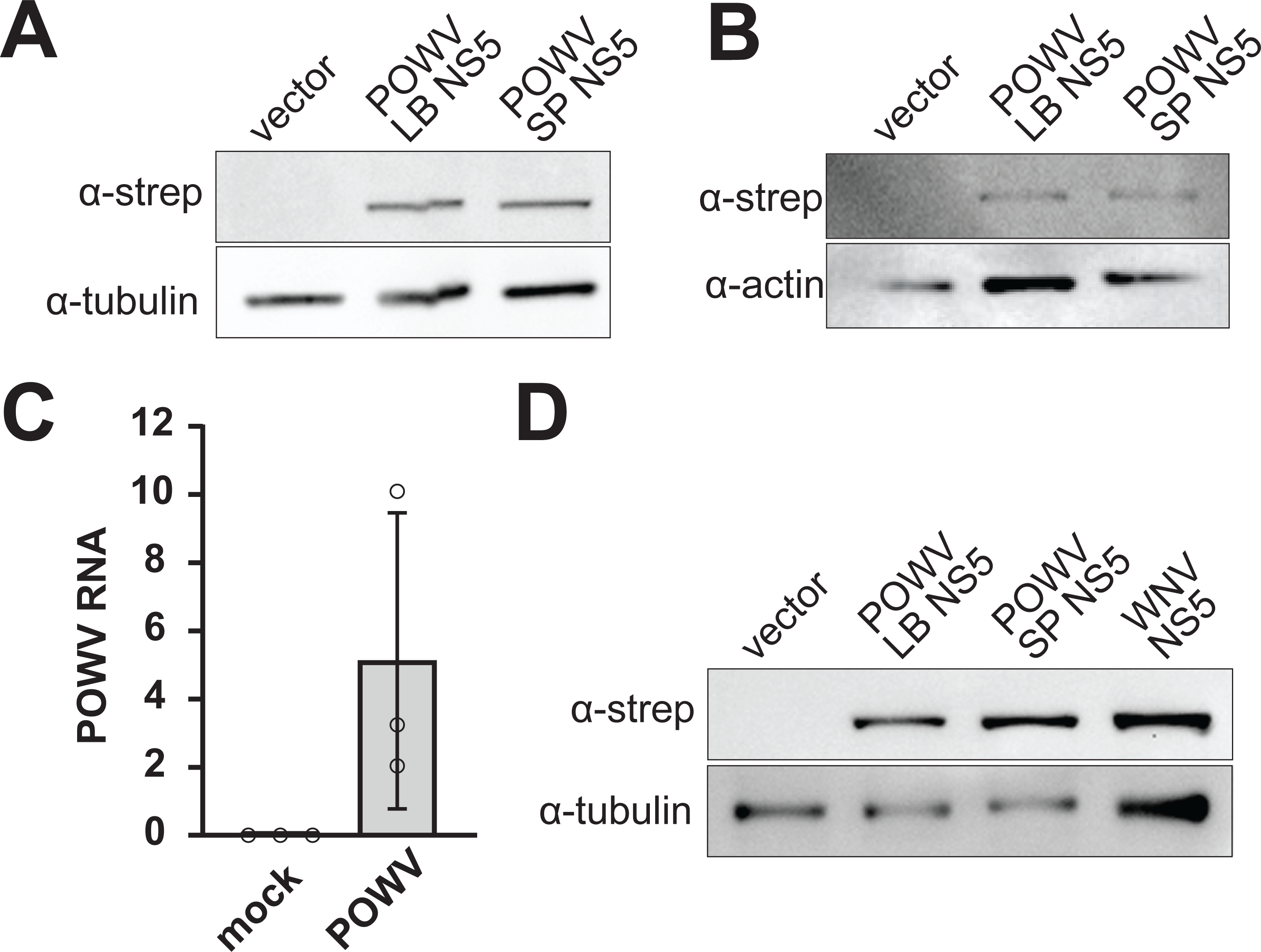
Lysates from HepG2 (**A**) and HaCaT (**B**) cells transduced with lentiviruses to stably express Strep-tagged POWV LB NS5, POWV SP NS5, or empty vector were analyzed via western blot with anti-Strep antibody. (**C**) HEK-Blue IL-12 cells were infected with POWV SP (MOI 1 for 48hr) and viral RNA measured by qRT-PCR. The mean ± SD is shown. (**D**) HEK-Blue IL-12 cells transduced with lentiviruses to stably express Strep-tagged POWV LB NS5, POWV SP NS5, WNV NS5, or empty vector were analyzed via western blot with anti-Strep antibody. Representative images shown.

## Supplementary Tables

**Table S1.** Host proteins interacting with POWV LB and POWV SP NS5 from HMC3 and HEK293T cells.

**Table S2.** Cloning primers, mutagenesis primers and gene blocks used in this study.

**Table S3.** Plasmids used in this study.

**Table S4.** Antibodies used in this study.

**Table S5.** qRT-PCR primers used in this study.

