## Supplementary Tables 2-5 for "Powassan Virus NS5 Antagonizes TYK2-Mediated Immune Signaling Pathways"

**Table S2: Cloning and mutagenesis primers and gene blocks**

| **Primers used to clone POWV genes:** | |
|---|---|
| POWV-LB-Capsid-F | GAGAAGGGGGCGGCCATGGTGACTACTTCTAAAGGAAAGG |
| POWV-LB-Capsid-R | AAACGGGCCCTCTAGCTAAGCCATCAAAGCCATTACC |
| POWV-LB-PrM-Env-F | CAGTGTGGTGGAATTATGGCCATGGCGACCTCC |
| POWV-LB-PrM-Env-R | CTCCCTCGAGCGGCCCGGCGTATACAGGCCCAAGG |
| POWV-LB-NS1-F | CAGTGTGGTGGAATTATGGACTATGGATGCGCAGTTG |
| POWV-LB-NS1-R | CTCCCTCGAGCGGCCCAGCCATTACCATAGATCTAACCAG |
| POWV-LB-NS2a-F | CAGTGTGGTGGAATTATGGACAACGGCGCGATGC |
| POWV-LB-NS2a-R | CTCCCTCGAGCGGCCCTCGTCTGCCATGGCCCC |
| POWV-LB-NS2b-F | CAGTGTGGTGGAATTATGAGTCTGAGTGAACCCC |
| POWV-LB-NS2b-R | CTCCCTCGAGCGGCCCTCTGCGTGTTGACGAGAG |
| POWV-LB-NS3-F | TACCGAGCTCGGATCATGACTGACCTTGTATTCTCAGGGC |
| POWV-LB-NS3-R | CTCCCTCGAGCGGCCCCCTGCGGCCAGAAGCGTA |
| POWV-LB-NS4a-F | CAGTGTGGTGGAATTATGAGTGCCATGGACATCTTCACTG |
| POWV-LB-NS4a-R | CTCCCTCGAGCGGCCCGCGTTGCTTTCCCGGTTC |
| POWV-LBNS4b-F | CAGTGTGGTGGAATTATGAATGAACTGGGCTATTTGGAGC |
| POWV-LB-NS4b-R | CTCCCTCGAGCGGCCCCCTTCTAGCCCCCTGAGTTC |
| POWV-LB-NS5-F | TTAAACTTAAGCTTGATGGGTGGAGCAGAGGGAAG |
| POWV-LB-NS5-R | CTCCCTCGAGCGGCCCGATTATTGAGCTCTCTAGCTTGAG |
| POWV-SP-NS5-F | TTAAACTTAAGCTTGATGGGTGGAGCAGAGGGAAG |
| POWV-SP-NS5-R | CTCCCTCGAGCGGCCCGATTATTGAGCTCTCTAGCTTGAG |

| Primers used to clone genes into pCDH lentiviral expression plasmid: | |
|---|---|
| Lenti-EV-Fwd | GCGATATCGATCCGGCGCGCCCCTGCAGGTTAATTAAG |
| Lenti-EV-Rev | CCGGATCGATATCGCTAGCTCTAGAATCTTCCCTCCATA |
| Lenti-WNV NS5-Fwd | CATAGAAGATTCTAGAATGGGTGGGGCAAAAGGAC |
| Lenti-WNV NS5-Rev | AGATCCTTCGCGGCCGCTTAAACGGGCCCCTTCTCG |
| Lenti-POWV-LB NS5-Fwd | CATAGAAGATTCTAGAATGGGTGGAGCAGAGGGAAG |
| Lenti-POWV-LB NS5-Rev | AGATCCTTCGCGGCCGCTTAAACGGGCCCCTTCTCG |
| Lenti-POWV-SP NS5-Fwd | CATAGAAGATTCTAGAATGGGTGGAGCAGAGGGAAG |
| Lenti-POWV-SP NS5-Rev | AGATCCTTCGCGGCCGCTTAAACGGGCCCCTTCTCG |
| **Primers used to generate POWV-LB NS5 mutants:** | |
| POWV-LB NS5-Mtase-Fwd | GTTTAAACTTAAGCTATGGGTGGAGCAGAGGGA |
| POWV-LB NS5-Mtase-Rev | TCCCTCGAGCGGCCCGGATGCCGCTGATCCGG |
| POWV-LB NS5-RdRp-Fwd | GTTTAAACTTAAGCTATGCTGATCAACGGAGTGG |
| POWV-LB NS5-RdRp-Rev | CTCCCTCGAGCGGCCCGATTATCGAGCTCTCTAGCTTGAGCTC |
| POWV-LB NS5-RdRp-453-Rev | CTCCCTCGAGCGGCCCCCACTTATTCTGTTCATCCGACCAT |
| POWV-LB NS5-RdRp-553-Rev | CTCCCTCGAGCGGCCCTAGCAGCCAGCCCAGATAGTT |
| POWV-LB NS5-RdRp-603-Rev | CTCCCTCGAGCGGCCCTTGGACTTTCATGTTTGTGATGGTGTTGAGGG |
| POWV-LB NS5-RdRp-703-Rev | CTCCCTCGAGCGGCCCCTTCATCACAAGTTCATGGAAGTGATGTG |
| POWV-LB NS5-RdRp-803-Rev | CTCCCTCGAGCGGCCCCAAAATCCAGACTTTGTTCCAGATGCG |
| POWV-LB NS5-5A-Fwd | CTGCTCTGGCAGCCGTAGCAACCTGGTTGAAGGAGCACG |
| POWV-LB NS5-5A-Rev | CGGCTGCCAGAGCAGGGGCCTGTGAGTCAGCTGGTCC |

| | |
|---|---|
| POWV-LB NS5-WNV NS5 swap-Fwd | CTGCTCTGGCAGCCGTAGCAACCTGGTTGAAGGAGCACG |
| POWV-LB NS5-WNV NS5 swap-Rev | CGGCTGCCAGAGCAGGGGCCTGTGAGTCAGCTGGTCC |
| POWV-LB-WNV chimera gBlock Amp-Fwd | CAACTCATCCGCATGATGGAAGGAGAAGGAGTCATAGGAC |
| POWV-LB-WNV chimera gBlock Amp-Rev | ACAATCATCTCCACTCACAAGCATCCTCCCTAGTC |

**Table S3: Plasmids used in this study**

| Construct name | Source |
|---|---|
| pcDNA4_TO-Strep-empty vector | This study |
| pcDNA4_TO-Strep-GFP | |
| pcDNA4_TO-Strep-WNV-NS5 | |
| pcDNA4_TO-POWV-Strep-Capsid | |
| pcDNA4_TO-POWV-LB PrM-Env-Strep | |
| pcDNA4_TO-POWV-LB NS1-Strep | |
| pcDNA4_TO-POWV-LB NS2a-Strep | |
| pcDNA4_TO-POWV-LB NS2b-3-Strep | |
| pcDNA4_TO-POWV-LB NS2b-Strep | |
| pcDNA4_TO-POWV-LB NS3-Strep | |
| pcDNA4_TO-POWV-LB NS4a-Strep | |
| pcDNA4_TO-POWV-LB NS4b-Strep | |
| pcDNA4_TO-POWV-LB NS5-Strep | |
| pcDNA4_TO-POWV-SP NS5-Strep | |
| pcDNA4_TO-POWV-LB NS5-Strep-MTase only | |
| pcDNA4_TO-POWV-LB NS5-Strep-RdRp only | |
| pcDNA4_TO-POWV-LB NS5-Strep-RdRp-453 | |
| pcDNA4_TO-POWV-LB NS5-Strep-RdRp-553 | |
| pcDNA4_TO-POWV-LB NS5-Strep-RdRp-603 | |
| pcDNA4_TO-POWV-LB NS5-Strep-RdRp-703 | |
| pcDNA4_TO-POWV-LB NS5-Strep-RdRp-803 | |

| | |
|---|---|
| pcDNA4_TO-POWV-LB NS5-Strep-5A | |
| pcDNA4_TO-POWV-LB NS5-Strep-WNV swap | |
| pDEST40-TYK2-DDAA-V5 | |
| pDEST40-TYK2-DDRR-V5 | |
| pDEST40-TYK2-V5 | Gracias et al., 2023 |
| pDEST40-TYK2-N-V5 | |
| pDEST40-TYK2-C-V5 | |
| pDEST40-TYK2-delta-KL-V5 | |
| pDEST40-TYK2-TK-V5 | |
| pDEST40-TYK2-delta-TK-V5 | |
| pDEST40-TYK2-KL-V5 | |
| pMD2.G | Addgene #12259 |
| psPAX2 | Addgene #12260 |
| pCDH_Str-Ii | Addgene #65313 |
| pGL4.45[luc2P/ISRE/Hygro] | Promega #E4141 |
| pNL1.1.TK[Nluc/TK] | Promega # N1501 |
| RIG-I 2Card | Gift from Dr. Michaela Gack (PMID: 26998762) |
| IFN-β luciferase reporter | Addgene #102597 |
| pCDH-empty vector-Strep | This study |
| pCDH-POWV-LB NS5-Strep | |
| pCDH-POWV-SP NS5-Strep | |
| pCDH-WNV NS5-Strep | |

**Table S4: Antibodies used in this study**

| Antibody | Source |
|---|---|
| Rabbit anti-TYK2 | Cell Signaling Technology #9312S |
| Rabbit anti-pY-TYK2 | Cell Signaling Technology #9321S |
| Mouse anti-IFNAR | Santa Cruz Biotechnology #sc-7391 |
| Rabbit anti-STAT1 | Cell Signaling Technology #14994S |
| Rabbit anti-pY-STAT1 | Cell Signaling Technology #7649L |
| Rabbit anti-STAT2 | Cell Signaling Technology #72604S |
| Rabbit anti-pY-STAT2 | Cell Signaling Technology #88410S |
| Mouse anti-Strep tag | Abcam #184224 |
| Rabbit anti-β-actin | CST #4967S |
| Mouse anti-α-tubulin | Sigma-Aldrich #T6199 |
| Mouse anti-LGTV E (cross-reacts with POWV E) | BEI Resources #NR-40316 |
| Rabbit anti-V5 tag | Bethyl # A190-120A |
| Mouse anti-V5 tag | Invitrogen #R960-25 |
| HRP goat anti-Rabbit IgG (H+L) | Invitrogen #G-21234 |
| HRP goat anti-Mouse IgG (H+L) | ThermoScientific #G-21040 |

**Table S5: qRT-PCR primers used in this study**

| Target | Forward primer | Reverse primer |
|---|---|---|
| 18S rRNA | GGCCCTGTAATTGGAATGAGTC | CCAAGATCCAACTACGAGCTT |
| POWV-SP | GCAGCACCATAGGTAGAATGT | CCACCCACTGAACCAAAGT |
| IFIT1 | GCGCTGGGTATGCGATCTC | CAGCCTGCCTTAGGGGAA |
| MX1 | GTTTCCGAAGTGGACATCGCA | CTGCACAGGTTGTTCTCAGC |
| RIG-I | CTGGACCCTACCTACATCCTG | GGCATCCAAAAAGCCACGG |
| SOCS1 | CACGCACTTCCGCACATTC | TAAGGGCGAAAAAGCAGTTCC |
